# Alternative host shapes transmission and life-history trait correlations in a multi-host plant pathogen

**DOI:** 10.1101/2022.06.17.496551

**Authors:** Hanna Susi

**Affiliations:** Research Centre for Ecological Change, Organismal and Evolutionary Biology Research Programme, Faculty of Biological and Environmental Sciences, University of Helsinki, P.O. Box 65, (Viikinkaari 1), 00014 University of Helsinki, Finland

**Keywords:** ecology, epidemiology, disease evolution, generalist pathogen, host-pathogen interaction, oomycete, landscape, trade-offs

## Abstract

Most pathogens are generalists capable of infecting multiple host species or strains. Trade-offs in performance among different hosts are expected to limit the evolution of generalism. Despite the commonness of generalism, the variation in infectivity, transmission, and trade-offs in performance among host species have rarely been studied in the wild. To understand the ecological and evolutionary drivers of multi-host pathogen infectivity and transmission potential, I studied disease severity, transmission dynamics and infectivity variation of downy mildew pathogen *Peronospora sparsa* on its three host plants *Rubus arcticus, R. chamaemorus*, and *R. saxatilis*. In a survey of 19 wild and cultivated sites of the three host species, disease severity varied by host species and by whether the focal host species grew in shared habitat with other host species. Generally, the presence of an alternative host resulted in lower disease severity. To understand how alternative host presence and plant diversity affect on transmission of the pathogen, I conducted a trap plant experiment. In contrast to the field survey observations, the presence of *R. saxatilis* was positively correlated with transmission to trap plants. To understand how resistance to *P. sparsa* varies among host species and genotypes, I conducted an inoculation experiment using 10 *P. sparsa* strains from different locations and 20 genotypes of the three host species. Significant variation in infectivity was found among host genotypes but not among host species. The pathogens strains originating form sites with *R. saxatilis* were less infective than those without. When trade-offs for infectivity were tested, high infectivity in one host species correlated with high infectivity in another host species. However, when pathogen transmission-related life-history correlations were tested, a positive correlation was found in *R. arcticus* but not in *R. saxatilis*. The results suggest that host resistance may shape pathogen life-history evolution with epidemiological consequences in a multi-host pathogen.

## Introduction

Most pathogens are generalists with the ability to infect multiple host species (Woolhouse et al., 2001). Generalism is common in all pathogen taxa e.g. malaria parasite *Plasmodium falciparum* (Hellgren et al., 2009), virus SARS-CoV-2 (Damas et al., 2020), white mold fungus *Sclerotinia sclerotiorum* (Bolton et al., 2006), and bacteria *Pseudomonas syringae* (Barrett et al., 2009). The ability to infect multiple host species gives a pathogen an advantage in spreading over wide areas and surviving in one host while another host goes locally extinct (Leggett et al., 2013). However, the pathogens may occur unnoticed in wild populations (Prendeville et al., 2012) and the disease symptoms may differ among host species (Saikkonen et al., 2016). These characteristics make generalist pathogens a challenge for their management and allow generalist pathogens to emerge and threaten the health of humans, crops and ecosystems (Maloney et al., 2005). Moreover, with increasing species introductions to new areas, attaining novel crops, and increased encounters for wild and cultivated species, the risks of emergence of multi-host pathogens have become more frequent (Jeger, 2022). Despite the well-recognized risks of generalist pathogens, it is surprising how little is known on the epidemiology and evolution of multi-host pathogens in the wild.

Despite the commonness of generalism, theory suggest that pathogens should favor specialism (Leggett et al., 2013). An expectation of a trade-off between high performance in one host as decreased performance in another has been suggested (Kawecki, 1994, Joshi & Thompson, 1995). Evidence for costs have been found in some cases whereas sometimes the evidence has been mixed (Leggett et al., 2013). An alternative cost of generalism has been sought between high infectivity among species vs performance within a host individual (Agudelo-Romero & Elena, 2008). This theory builds on the hypothesis that if the pathogen can encounter many hosts and infect them it does not need to reach high prevalence within host individuala to persist (Combes, 1997). However, Hellgren et al (Hellgren et al., 2009) showed that the pathogens with high infectivity among host species were also capable of reaching high prevalence within the host individuals. Similarly, in a study of multi-host fungal pathogen *Microbotryum*, Bruns et al. (2021) found that the pathogens with high intraspecific infectivity were also capable of high sporulation in their shared host species. On the other hand, infecting novel host species provides an escape from the host resistance mechanisms for the pathogen (Thines, 2019, Barrett et al., 2009). It is suggested that life-history trait correlations may be altered in a host where the pathogen has not evolved (Thines, 2019) leading to a situation where the pathogen persists within a host with low transmission and low harm for the host (Saikkonen et al., 2016).

Additional important drivers for evolution and epidemiology of multi-host pathogens are community context and environmental variability (Haas et al., 2011, Gilligan & van den Bosch, 2008, Woolhouse et al., 2001). In an environment where alternative hosts are abundantly available increased virulence is expected whereas lower virulence is expected when a single host species is present (Frank, 1992). In addition to availability of susceptible host, the overall community diversity shapes transmission of multi-host pathogens (Haas et al., 2011). High diversity in the community where the host is embedded is expected to dilute disease transmission (i.e. dilution effect (Rohr et al., 2020)). An alternative scenario is amplification, especially in the case of multi-host pathogens, in which an increase in the number of competent hosts accelerates disease spread (Begon, 2008). However, asymmetry in the host compatibility adds yet another layer of complexity to disease transmission within the community context (Pedersen & Fenton, 2007, Haas et al., 2011). Resistance variation within and among species may thus have a profound effect on epidemiology and emergence of multi-host pathogens (Woolhouse et al., 2001).

When novel crop species are taken to cultivation or a crop is cultivated in new areas, generalist pathogens existing in the same or alternative hosts in the wild may generate pathogen spill over leading to severe crop losses (Jeger, 2022). Understanding the host ranges, epidemiology and transmission dynamics of local pathogen species is essential to prevent emergence of pathogens in changing crop selection (Jeger, 2022). *Rubus arcticus*, the arctic bramble, a berry crop with high value yields has been taken to cultivation since 1970’s in Finland. However, a downy mildew pathogen disease caused by oomycete *Peronospora sparsa*, also known as ‘dryberry’ disease, has caused devastating losses in the cultivation of the berry (Koponen et al., 2000). The downy mildew disease outbreaks lead to losses of up to 50% in marketable yield by causing drying of the developing berries (Koponen et al., 2000). The pathogen favors cool and moist climate and spreads via air and water droplets. Current control methods against the pathogen rely on fungicides as there are no resistant cultivars available (Kostamo et al., 2015, Parikka et al., 2016). *Peronospora sparsa* is a multi-host pathogen (Hukkanen et al., 2006) and in addition to wild *R. arcticus* its other potential host species cloudberry, *Rubus chamaemorus*, and stone bramble, *Rubus saxatilis*, grow commonly in forests and roadsides adjacent to the plantations making pathogen transmission between fields and wild plants a challenge for disease management. The pathogen has been reported to occur commonly in wild populations of *R. arcticu*s whereas its infectivity to *R. chamaemorus* has only been shown in laboratory trials and herbaria samples (Koponen et al., 2000).Previous attempts to detect the pathogen on *R. saxatilis* growing adjacent to arctic bramble plantations have not found it (Koponen et al., 2000). Currently, the pathogen’s ecology and adaptation to its host species is poorly understood.

Here, I studied the host range, transmission dynamics, and life-history correlations of the oomycete multi-host pathogen downy mildew pathogen *Peronospora sparsa* on its three *Rosacean* host species *Rubus arcticus* arctic bramble*, R. chamaemorus* cloudberry, and *R. saxatilis* stone bramble by conducting a field sampling, a trap plant experiment, and a laboratory inoculation experiment. To understand the pathogen’s natural host range and prevalence in the three host species, I detected the pathogen from plant samples collected from 19 sites using a PCR test. Transmission dynamics were observed by placing *R. arcticus* trap plants in 20 *R. arcticus* sites with varying presence of *R. saxatilis* and *R. chamaemorus*. To understand *P. sparsa* performance in the three host species, a laboratory inoculation trial using 10 *P. sparsa* strains originating from wild sites was set up. In all three data sets, I used latitude as a determinant of pathogen background and site environment. Latitude is expected to influence host-pathogen interactions (Zvereva & Kozlov, 2021, Magnusson et al., 2020, Liu et al., 2020) and increased disease pressure is predicted in higher latitudes (Bebber et al., 2013). More specifically, the aim of the study was to test 1) Does *P. sparsa* infect *R. saxatilis* and *R. chamaemorus* in the wild, and how common the symptoms on the three species in wild versus cultivated populations? 2) Does transmission of *P. sparsa* vary by plant diversity and host abundance? I expect to find dilution effect by plant diversity (Liu et al., 2020) and increase in transmission by host abundance (Frank, 1992, Woolhouse et al., 2001). 3) Does performance of *P. sparsa* in the three host species vary in a laboratory inoculation experiment and are there trade-offs between pathogen life-history traits? I expect that high performance in one host has a cost as poorer performance in another (Kawecki & Ebert, 2004). 4) Does abundance of *R. saxatilis* affect pathogen infectivity and transmission potential in a laboratory experiment? I expect infectivity to increase in pathogen strains originating from populations with alternative host (Woolhouse et al., 2001).

*Peronospora sparsa* infection was confirmed by PCR test in all three *Rubus* species in the wild. *Peronospora sparsa* occurrence and symptom prevalence varied among the host species being highest in *R. chamaemorus* and lowest in *R. saxatilis*. In the transmission experiment, coverage of *R. saxatilis* increased *P. sparsa* infection occurrence and severity. In sites with *R. saxatilis*, plant diversity diluted transmission to trap plants. In the laboratory experiment, all 10 pathogen strains could infect all three host species while most of the variation in infectivity was among host genotypes within host species. High infectivity in one host species was correlated with high infectivity in another species. However, when trade-offs among life-history traits within species were tested for, fast sporulation correlated with abundant sporulation in *R. arcticus* while in *R. chamaemorus* and *R. saxatilis* the correlation did not exist. In addition, pathogen strain infectivity correlated negatively with prevalence of *R. saxatilis* in the site of origin. Jointly these results suggest that while alternative host abundance increased transmission potential, there are evolutionary constrains limiting the pathogen performance in alternative hosts.

## Material and methods

### The hosts and the pathogen

*Rubus arcticus* L, arctic bramble, is a perennial diploid plant belonging to *Rosacea*. Its native distribution spans over subarctic Eurasia and Asia to North America. The plant is an obligate out crosser and spreads vegetatively via rhizomes (Tammisola, 1988). *R. arcticus* is cultivated for commercial use in Finland but it’s yields have been severely damaged by downy mildew disease caused by *Peronospora sparsa* (Koponen et al., 2000). *Rubus chamaemorus*, cloudberry, is a perennial dioecious wild plant native to northern hemisphere. It occurs in mountainous areas and moorlands. Despite some attempts to cultivate the plant for its berries and leaves, *R. chamaemorus* is not widely cultivated. *Rubus saxatilis*, stone bramble, is a perennial plant distributed in temperate regions in Eurasia commonly occurring in forests and field sides. The plant is not commercially cultivated. *Rubus arcticus* and *R. saxatilis* are phylogenetically close relatives belonging to *Cylactis* subgenus whereas *R. chamaemorus* belongs to *Chamaemorus* subgenus (Sobczyk, 2018).

*Peronospora sparsa* is an oomycete pathogen infecting *R. arcticus* in Nordic countries (Koponen et al., 2000). The pathogen can be visually observed as purple dark lesions on the leaves and dry, malformed berries. The pathogen is an obligate biotroph (Thines & Choi, 2016) and several asexual cycles during the growing season spread the pathogen (Kostamo et al., 2015). Asexual spores are spread by air and water and germination occurs in moist and cool conditions (Aegerter et al., 2003). *Peronospora sparsa* is divided in two subspecies one infecting genus *Rosa* and another infecting genus *Rubus* (Thines & Choi, 2016). Laboratory inoculations and herbaria samples have shown the pathogen to infect *R. arcticus* and *R. chamaemorus* in the Nordic countries (Koponen et al., 2000). Despite attempts to survey *R. saxatilis* populations adjacent to berry farms there are no records on the pathogen in *R. saxatilis* (Koponen et al., 2000).

### Survey and detection of *P. sparsa* in cultivations and wild sites

To investigate the host range of *P. sparsa* in naturally occurring *Rubus* genera in Finland, I surveyed 20 sites (14 natural sites and 6 plantations; Supplementary Table 1) across Finland in late August 2019. At each site, *P. sparsa* infection prevalence in 10-30 plants of each *Rubus* genera was observed. In addition, host plant population size was estimated as square meters covered by the host species in each site. Infection prevalence was estimated as the percentage of infected hosts with visible symptoms. To confirm that *P. sparsa* is the causal agent of the disease symptoms, 2 cm^2^ piece of symptomatic leaf was collected into a microtube and stored in −80°C until DNA extraction. Samples were collected from 10 plants from each species from each site.

In laboratory, DNA was extracted from the samples using CTAB method as in (Lodhi et al., 1994). The yield and quality of DNA was measured using Nanodrop. PCR reaction was set up using primers P1 and P2 (Hukkanen et al., 2006) in following conditions: 10 μl PCR mix included 0.5 volume GoTaq^®^ Green Master Mix (Promega), 1 μl primer P1(10μM), 1 μl primer P2(10μM), 3 μl MilliQ water, and 1 μl template. The reaction cycle was 92°C 2 minutes, 30 cycles 92°C 30 seconds, 56°C 30 seconds and 72°C 30 seconds with final extension in 72°C 5 minutes. The PCR product was subjected to gel electrophoresis and visualized using BioRad.

### Transmission experiment

Plant material for transmission experiment was obtained by cloning small plants from rhizomes in insect-free greenhouse in 16:8 day: night conditions in 17°± 2°C for four months. The clones were separated from the plant as terminal buds of the rhizome and placed in to separate 8 × 8 cm pots filled with mixture of 25 % potting soil, 25 % lightweight expanded clay aggregate, and 50 % peat. After cloning, the dormant buds were kept in +4 °C for 4 months until the experiment. Thus, all plants were in the same early stages of development when placed into the field sites. Three pathogen free genotypes were used in the experiment, commercial cultivar *Pima*, and wild genotypes G12 and G13 originating from Muuruvesi and Viinikka, respectively. To understand how presence of alternative hosts and the diversity of surrounding plant diversity may affect *P. sparsa* transmission and within host infection prevalence, I set up a field experiment using 600 plants distributed to 20 sites (30 plants per site). The sites were selected so that there were sites in which only *R. arcticus* (10 sites) was present and sites where either *R. chamaemorus* (1 site) or *R. saxatilis* (9 sites) was present in addition to *R. arcticus*. To understand how latitude affects disease prevalence, transmission dynamics, and pathogen performance, the sites were selected along a 1200 km latitudinal gradient. FinBIF Database (FinBIF, https://laji.fi/taxon/MX.mesimarja (accessed 14.06.2021)) was used in site selection and in determining the prevalence of *Rubus* species. The experiment was set up in mid-June 2021. At each site, *R. arcticus* trap plants consisting of three genotypes were placed randomly in 0.5-meter distance of *Rubus* plants in their pots. The plants were left to the sites for seven weeks after which they were revisited, and the infection status and prevalence was observed in each plant. To evaluate the association between *P. sparsa* transmission and plant diversity, the vascular plant species were identified in 3-6 one square meter vegetation plots (number of plots depended on the area of the site) in each site. The coverage of each species was assessed in each plot and the prevalence of each species on site level was calculated as the sum of its coverage within the vegetation plots in the site. Shannon’s index of diversity (Shannon & Weaver, 1949) in each site was calculated based on the plant species and their prevalence in the site as:

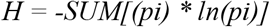

where SUM is summation, and pi is the prevalence of a plant species as the summed occurrence (0 = species not found in the given site; 1 = species found in the given site) of species/total number of vegetation plots within each site.

### Inoculation experiment

To confirm the host range of *P. sparsa* and to compare its performance and life-history trait correlations among *R. arcticus, R. saxatilis*, and *R. chamaemorus* an inoculation experiment was set up. Ten *P. sparsa* strains (Supplementary Table 2) originating from different sites were used in the experiment. Infected *R. arcticus* leaves were collected from the sites and placed on Petri dishes and brought to laboratory. In laboratory, the spores of the pathogen were inoculated onto young, detached leaves of susceptible *R. arcticus* plants on Petri dishes. The inoculated leaves were maintained in 16:8 day: night conditions in 17°±2°C for two weeks after which they were transferred onto new leaves. The plant material for the inoculation experiment consisted of 20 host genotypes (11 *R. arcticus* genotypes, 4 *R. chamaemorus* genotypes and 5 *R. saxatilis* genotypes; Supplementary Table 2) that were inoculated with the ten *P. sparsa* strains making altogether 200 host genotype–pathogen genotype combinations. Each combination was replicated three times leading to 600 inoculations in total. Detached leaves placed on moist filter paper on Petri dishes were inoculated with conidial spores from 1 cm^2^ sized lesions by evenly brushing the spores on a leaf with moist paint brush. Pathogen development was observed daily under a dissecting microscope starting at seventh day post-inoculation (DPI) until day 21 DPI. Infectivity was recorded as 0 = no infection and 1 = infection (sporulation observed). I measured two components of pathogen transmission potential: time to pathogen sporulation and pathogen lesion development at day 21 DPI. The first day when spores were observed was considered as the day of sporulation. Pathogen lesion development was measured on a scale from 0 to 4: 0 = no mycelium, 1 = only mycelium; 2 = mycelium and sparce sporulation visible only through microscope; 3 = abundant sporulation, colony size < 0.5 cm^2^; and 4 = abundant sporulation and colony size > 0.5 cm^2^ (Bevan et al., 1993).

### Statistical analyses

All statistical analyses were performed in SAS. To understand whether there was variation in symptom occurrence and prevalence across *Rubus* sites, I ran a set of statistical modeld on the survey data collected in 2019 using Generalized linear model framework in SAS GLIMMIX (SAS Institute). First, I analyzed whether the three *Rubus* species differ in their Symptom prevalence measured as the percentage of diseased plants in each site. Symptom prevalence measured as the percentage of diseased plants in each site was used as continuous response variable and plant population size measured as the coverage of the focal host plant in square meters were used as continuous explanatory variables. Focal host species, alternative host species and cultivated vs wild were used as class explanatory variables. Secondly, I fit another model to understand the drivers of symptom prevalence measured as the proportion of infected leaf area in each surveyed plant as response variable. In this model, host population size measured as the coverage of focal host species in square meters was used as continuous explanatory variable. Focal host species, alternative host species and cultivated vs wild were used as class explanatory variables. In both models, Gamma distribution of error was used. In these analyses, the interactions between model variables were tested and the significant interactions with best Akaike information criterion was used in the model selection.

To explore the drivers of transmission of *P. sparsa* in sites with varying prevalence of *Rubus* species and plant diversity, two Generalized linear models were set up. The data collected in the trap plant experiment in 2021 was used in these analyses. In the first model, I used the infection status of each trap plant as a binary response variable. Shannon diversity of plants, *R. arcticus* coverage, *R. saxatilis* coverage, *R. chamaemorus* coverage, and latitude were used as continuous explanatory variables. Site and trap plant genotype were used as class explanatory variables. Binomial distribution of error was assumed. To understand the drivers of the within host transmission of the pathogen, I fit another model with similar explanatory structure but used the proportion of infected leaves within trap plant as continuous response variable. In this model, a Gamma distribution of error was assumed. In these models, the interactions between variables were tested and the significant interactions with best Akaike information criterion was used in the model selection.

To test whether there were correlations among site variables in the transplant experiment (Shannon diversity of plants, *R. arcticus* coverage, *R. saxatilis* coverage, *R. chamaemorus* coverage, and latitude), a set of nine regression analyses was conducted in SAS Proc REG (SAS Institute) by regressing each variable against each other variable separately.

To assess the effects of pathogen genotype, host species and genotype on pathogen performance, three models were set up. First, to test the variation in infectivity of the pathogen strains, a model using the infection outcome (0 = no infection; 1 = infection) as binary response variable was set up. Pathogen strain, host species, and host genotype nested under host species were used as class explanatory variables. Binomial distribution of error was assumed. The second model with similar structure was set up to assess the speed of sporulation. In this model, I used speed to sporulation measured as the day ofsporulation deducted from 21 days (the end of the experiment) as a continuous response variable. A Gamma distribution of error was assumed. The third model with similar structure was fitted to test the variation in *P. sparsa* lesion development measured on a scaled from 0 to 4: 0 = no mycelium, 1 = only mycelium; 2 = mycelium and sparce sporulation visible only through microscope; 3 = abundant sporulation, colony size < 0.5 cm^2^; and 4 = abundant sporulation and colony size > 0.5 cm^2^ (Bevan et al., 1993). In this model, a Lognormal distribution of error was assumed. In the three models, the significance of interactions between latitude of pathogen origin and host species as well as host genotype nested under host species were tested to understand the specificity of the interaction between the host and the pathogen. To test whether the host plant combination at the site of origin influenced the pathogen performance, another set of three models with a similar response and explanatory model structure was run. In these analyses, in these analyses, *R. saxatilis* ja *R. arcticus* population sizes were added to the explanatory variables. In these models, the significance of interactions between *R. arcticus* and *R. saxatilis* coverage, and the other variables was tested.

Lastly, I set up four models to test whether there are trade-offs between pathogen life history traits and whether host species affects the potential life-history trait correlations. First, to test whether high infectivity in one host species has a cost as reduced infectivity in other host species, three models in PROC REG in SAS (SAS Institute) using the mean infectivity across each *P. sparsa* strain on each host species as response variable were run. Secondly, to understand whether fast sporulation corresponds with sporulation abundance at the end of the experiment, and whether host species influences the potential correlation, I ran a model in PROC GLIMMIX in SAS (SAS Institute) using the mean sporulation stage of the pathogen on each host genotype as a continuous response variable. Speed to sporulation (the day of first sporulation deducted from 21 days) was used as continuous explanatory variable. Pathogen strain, host species, and host genotype nested under host species were used a class explanatory variable. Interactions between host species, host genotype nested under host species, pathogen strain, and speed to sporulation were tested, and only statistically significant interactions were kept in the model. A Gaussian distribution of errors was assumed.

## Results

To understand the prevalence and host range of *P. sparsa* in wild and cultivated *Rubus* species, a field survey in 20 sites was performed. PCR detections on symptomatic plants of three species *R. arcticus, R. saxatilis* and *R. chamaemorus* were done. In addition to wild and cultivated *R. arcticus* sites, *P. sparsa* was detected in two novel host plants in the wild, *R. chamaemorus* and *R. saxatilis* by PCR. In the infected sites, high symptom occurrence was found in *R. arcticus, R. chamaemorus* and *R. saxatilis* (Fig. 1). The three host plants differed in symptom occurrence, *R. chamaemorus* having the highest mean symptom occurrence of 64%, measured as the percentage of plants showing symptoms in a site, *R. arcticus* mean symptom occurrence being 58% and *R. saxatilis* 36% (Table 1; Fig. 1a). Difference in symptom prevalence (as proportion of symptomatic leaf area) was pronounced between *R. chamaemorus* (19.6%), and the two other hosts *R. arcticus* (6.8%) and in *R. saxatilis* (3.6%; Table 1; Fig. 1b). However, variation among the hosts differed among sites (significant interaction site × species; Table 1; Fig 1b). In cultivated sites, symptomatic leaf area was smaller than in wild sites (Table 1; Fig. 1d) whereas symptom occurrence did not differ between wild and cultivated sites (Table 1). In 11 sites two host species grow together, and the presence and identity of the alternative host was linked to the symptom prevalence of the focal host (Table 1; Fig. 1c). The presence of *R. saxatilis* was associated with lower symptom prevalence and absence of alternative host was associated with lower symptom prevalence (Table 1; Fig. 1c). However, the effect of alternative host species was dependent on the focal host species (significant interaction: focal host species × alternative host species; Table 1; Fig. 1c).

**Figure 1.**
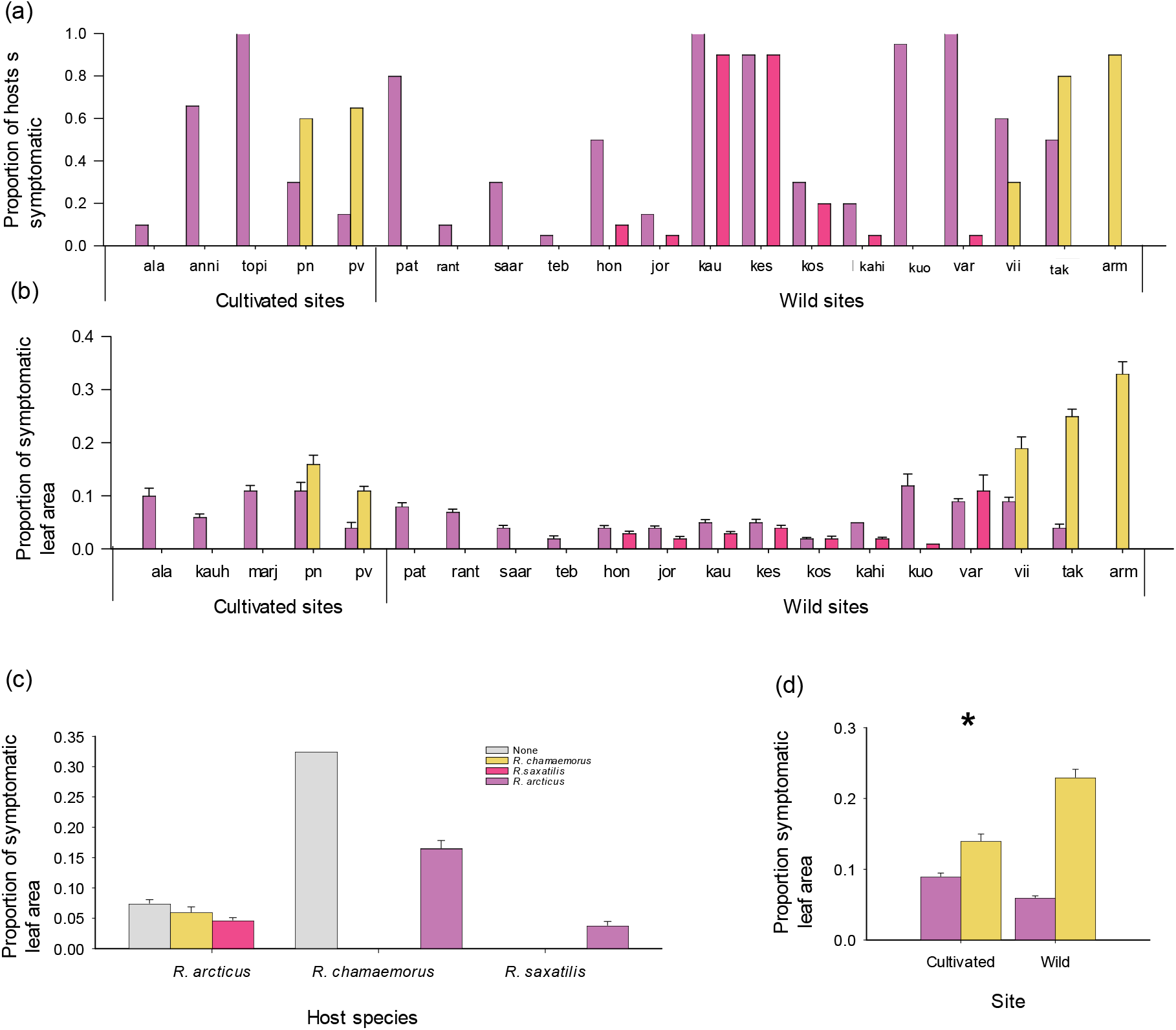
Variation in *Peronospora sparsa* symptoms in three *Rubus* hosts in the wild. The variation in *Peronospora sparsa* (a) symptom occurrence and (b) prevalence (as proportion of infected leaf area) on *Rubus arcticus, R. chamaemorus*, and *R. saxatilis* in 20 sites in 2019. (c) The variation in symptom prevalence in *Rubus* growing in habitats with and without other *P. sparsa* hosts. (d) The variation in symptom prevalence in wild and cultivated hosts.

**Table 1.** The factors affecting *Peronospora sparsa* symptom prevalence and symptom occurrence in 20 sites of *Rubus arcticus, R. chamaemorus*, and R. *saxatilis* surveyed in Finland in 2019. The results were analyzed by Generalized linear models. Statistically significant (*P* < 0.05) results are shown in bold.

| Effect | Symptom prevalence |  |  | Symptom occurrence |  |  |
| --- | --- | --- | --- | --- | --- | --- |
|  | *D.F.* | *F* | *P* | *D.F.* | *F* | *P* |
| Site | 16, 421 | 6.36 | <.0001 | 20, 9 | 13.32 | 0.0002 |
| Focal host coverage | 1, 421 | 4.88 | 0.0276 | 1, 9 | 19.68 | 0.0016 |
| Cultivated vs wild | 1, 421 | 5.72 | 0.0172 | 1, 9 | 1.26 | 0.2908 |
| Focal host species | 2, 421 | 13.44 | <.0001 | 2, 9 | 29.85 | 0.0001 |
| Alternative host species | 3, 421 | 5.14 | 0.0017 |  |  |  |
| Focal host species × alternative host species | 1, 421 | 6.91 | 0.0089 |  |  |  |

To understand how variation in host coverage and plant diversity shape transmission dynamics of *P. sparsa* in the wild, a trap plant experiment was set up in 21 sites along a 1200 km latitudinal gradient (Supplementary Figure S3) by placing potted *R. arcticus* plants in each site for seven weeks. When the sites were revisited, altogether 78% (466) plants had survived in the sites (20-100% plants survived within each site). I found that the plant size measured as number of leaves, genotype, and *R. saxatilis* coverage caused the largest variation in infection occurrence of the plants (Table 2; Fig 2a). There was no effect of latitude on the infection occurrence (Table 2), but sites varied with between 55-100% trap plants diseased (Supplementary Table 2; Fig 2a). Overall, the highest infection occurrence was observed in three sites, Korretoja, Hongikonluoma, and Säärenperä, where all plants became infected and lowest in Pahkalahti where only 55% of the plants became infected (Supplementary Table 2). Trap plant genotype G12 had the highest infection occurrence whereas in G13 it was the lowest. Plant size correlated positively with infection occurrence. Host plant combination in the study sites was linked to *P. sparsa* transmission to trap plants. *Rubus saxatilis* coverage increased infection occurrence but there was no significant effect of *R. arcticus* on infection occurrence on the trap plants (Table 2; Fig. 2a). Shannon diversity had no direct effect on infection occurrence (Table 2). However, *R. saxatilis* coverage altered the effect of Shannon diversity so that the negative correlation between Shannon diversity and infection occurrence became more pronounced in the sites with *R. saxatilis* (significant interaction *R. saxatilis* coverage × Shannon diversity; Table 2; Figure 2c). When infection prevalence (proportion of infected leaves within a plant) was analyzed, I found that *R. arcticus* and *R. saxatilis* coverage correlated positively with high infection prevalence (Table 2; Fig. 2b). The effect of plant diversity (Shannon’s) was again mediated by *R. saxatilis* coverage (significant interaction *R. saxatilis* coverage × Shannon diversity; Table 2; Figure 2d). In the sites with *R. saxatilis* presence, plant diversity correlated negatively with infection prevalence whereas in the sites without *R. saxatilis* the opposite was observed. There was a small but significant negative correlation between latitude and infection prevalence (Table 2). Large plants became less infected than small plants (Table 2), and trap plant genotypes varied in their infection prevalence (Table 2; Supplementary Fig. S2). The regression analysis on the site variables indicated that there were no significant correlations among them (Supplementary Table S4).

**Figure 2.**
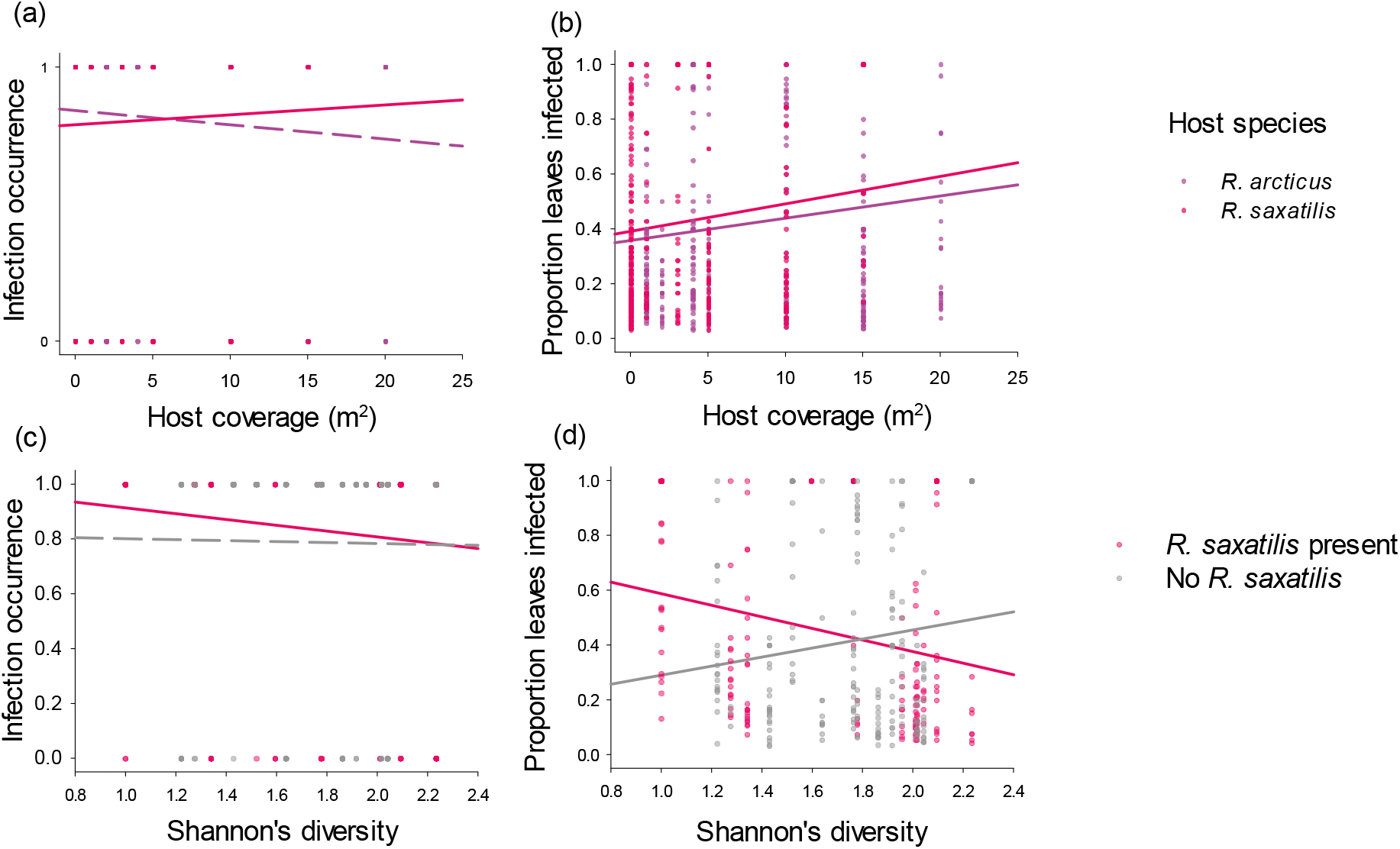
Variation in *Peronospora sparsa* transmission to *Rubus arcticus* trap plants in 20 sites with varying alternative host abundance. The correlation between *Rubus arcticus* and *R. saxatilis* coverage and (a) infection occurrence and (b) prevalence (as proportion of infected leaves) in trap plants (n = 493). The effect of *R. saxatilis* presence on the correlation between plant diversity (*Shannon’s*) and *P. sparsa* (c) infection occurrence and (d) infection prevalence.

**Table 2.** The factors affecting transmission of *Peronospora sparsa* to *Rubus arcticus* trap plants in 21 sites in Finland. The results were analyzed by Generalized linear models. Statistically significant (*P* < 0.05) results are shown in bold.

| Effect | Infection occurrence |  |  | Infection prevalence |  |  |
| --- | --- | --- | --- | --- | --- | --- |
|  | *D.F.* | *F* | *P* | *D.F.* | *F* | *P* |
| Latitude | 1, 445 | 0.11 | 0.7416 | 1, 359 | 4.9 | 0.0275 |
| Plant size | 1, 445 | 24.52 | <.0001 | 1, 359 | 66.8 | <.0001 |
| Plant genotype | 2, 445 | 5.88 | 0.003 | 2, 363 | 8.78 | 0.0002 |
| *R. saxatilis* coverage | 1, 445 | 8.03 | 0.0048 | 1, 359 | 12.93 | 0.0004 |
| *R. arcticus* coverage | 1, 445 | 7.77 | 0.0055 | 1, 359 | 1.69 | 0.1947 |
| Shannon's diversity | 1, 445 | 1.04 | 0.3083 | 1, 359 | 0.2 | 0.6579 |
| *R. saxatilis* coverage × Shannon's diversity | 1, 445 | 7.26 | 0.0073 | 1, 359 | 10.63 | 0.0012 |

To disentangle host susceptibility, pathogen infectivity and transmission potential in the interaction among *Peronospora sparsa* isolates originating from a latitudinal gradient and its three *Rubus* host species, an inoculation experiment was set up. When pathogen infectivity (i.e., host susceptibility) was tested, the northern pathogen strains had higher infectivity than the southern ones (Table 3; Fig. 3a). Generalism was common among the studied pathogen strains as six of the ten *P. sparsa* strains were able to infect all three host species (Fig. 3a). None of the *P. sparsa* stains was able to infect two of the *R. chamaemorus* genotypes (Fig. 3b). The infectivity across the plant genotypes used in the experiment varied between 35-75% (Fig. 3a). Host genotypes within host species differed significantly in their susceptibility but host species did not explain a significant amount of variation (Table 3; Fig. 3b). The susceptibility of the plant genotypes ranged from 0% to 100% of the pathogen strains used (Fig. 3b). When the factors driving transmission potential i.e., speed to sporulation and sporulation abundance were analyzed, there were significant differences between genotypes within species but neither the latitude of pathogen origin nor host species had significant effect on these life-history traits (Table 3; Fig. 3c-f). In all three models, the interaction between latitude and host species as well as genotype of the host nested under host species were tested but as they were nonsignificant, they were excluded from the final models.

**Figure 3.**
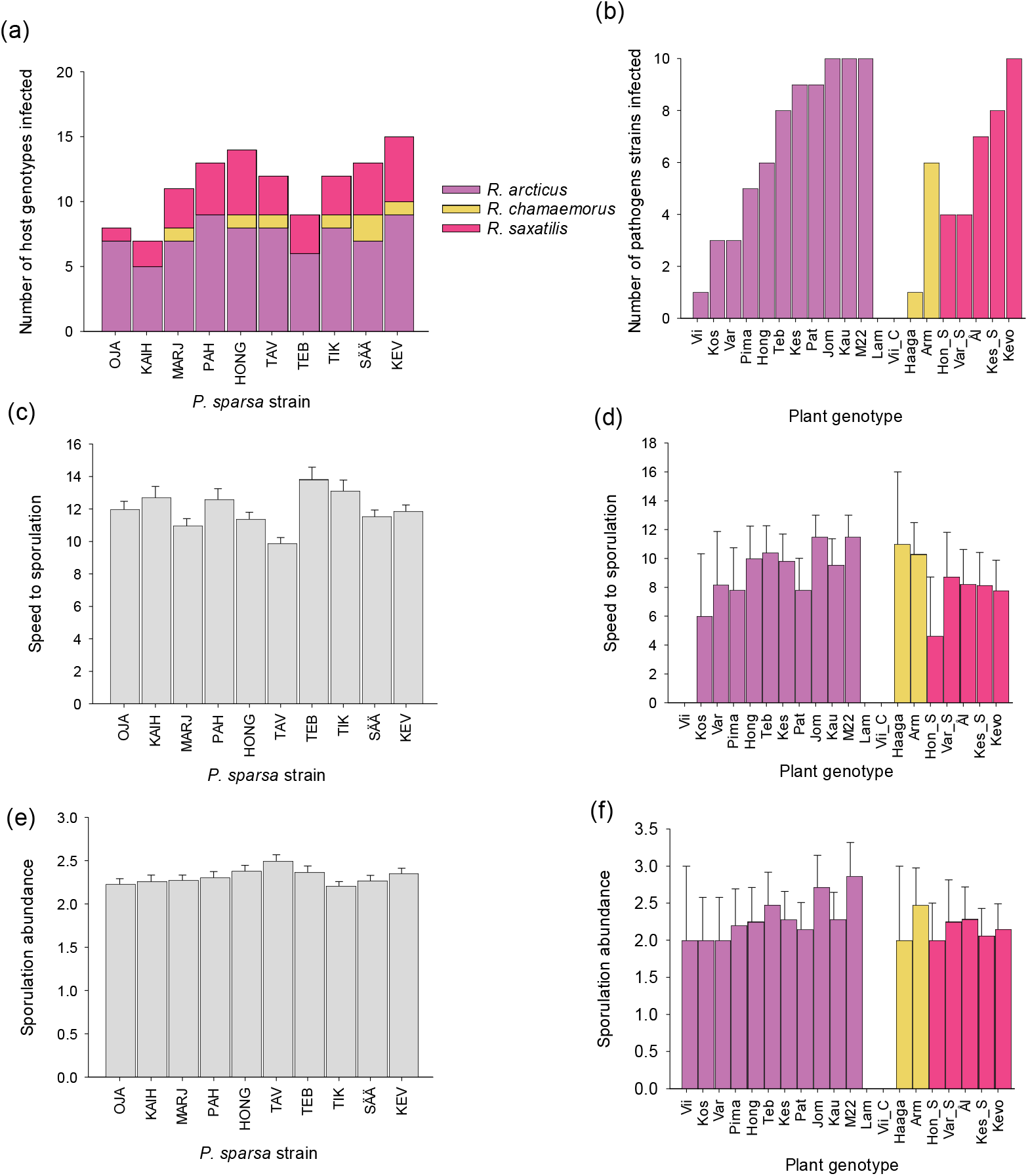
Variation in infectivity and transmission potential in an inoculation experiment on 10 *Peronospora sparsa* strains and 11 *Rubus arcticus*, 4 *R. chamaemorus* and 5 *R. saxatilis* genotypes. (a) Variation among *P. sparsa* strains (shown in a latitudinal order) in their ability to infect host genotypes. (b) The variation among *Rubus* genotypes in the infectivity of *P. sparsa*. The variation in speed to first sporulation (21 days – the day when first sporulation was observed in microscopy) among (c) *P. sparsa* strains and (d) *Rubus* genotypes. The variation in sporulation abundance (e) *P. sparsa* strains and (f) *Rubus* genotypes.

**Table 3.** The results from a laboratory inoculation experiment on 10 *Peronospora sparsa* strains originating from a latitudinal gradient and 20 plant genotypes belonging to *Rubus arcticus, R. chamaemorus* and *R. saxatilis*. The results were analyzed by Generalized linear models. Statistically significant (*P* < 0.05) results are shown in bold.

| Effect | Infectivity |  |  | Speed to sporulation |  |  | Sporulation abundance |  |  | Sporulation abundance |  |  |
| --- | --- | --- | --- | --- | --- | --- | --- | --- | --- | --- | --- | --- |
|  | D.F. | F | P | D.F. | F | P | D.F. | F | P | D.F. | F | P |
| Latitude | 1, 179 | 8.16 | 0.0048 | 1, 95 | 0.43 | 0.5146 | 1, 95 | 1.13 | 0.2904 | 1, 92 | 0.24 | 0.6263 |
| Species | 2, 179 | 0 | 0.9974 | 2, 95 | 2.43 | 0.0935 | 2, 95 | 1.45 | 0.2397 | 2, 92 | 4.5 | 0.0137 |
| Genotype within species | 17, 179 | 1.73 | 0.0415 | 15, 95 | 2.11 | 0.0154 | 15, 95 | 3.07 | 0.0005 | 15, 92 | 1.98 | 0.0254 |
| Speed to sporulation |  |  |  |  |  |  |  |  |  | 1, 92 | 4.25 | 0.0422 |
| Speed to sporulation × species |  |  |  |  |  |  |  |  |  | 2, 92 | 3.96 | 0.0224 |
| Effect | Infectivity |  |  | Speed to sporulation |  |  | Sporulation abundance |  |  |  |  |  |
|  | D.F. | F | P | D.F. | F | P | D.F. | F | P |  |  |  |
| Latitude | 1, 177 | 5.81 | 0.0169 | 1, 93 | 0.22 | 0.6431 | 1, 93 | 0.43 | 0.5152 |  |  |  |
| Species | 2, 177 | 0 | 0.9989 | 2, 93 | 1.64 | 0.1994 | 2, 93 | 1.55 | 0.2173 |  |  |  |
| Genotype within species | 17, 177 | 1.74 | 0.0403 | 15, 93 | 1.6 | 0.0884 | 15, 93 | 3.07 | 0.0005 |  |  |  |
| *R. arcticus* coverage | 1, 177 | 0.52 | 0.4709 | 1, 93 | 1.95 | 0.166 | 1, 93 | 0.07 | 0.7964 |  |  |  |
| *R. saxatilis* coverage | 1, 177 | 5.25 | 0.0231 | 1, 93 | 0.4 | 0.5274 | 1, 93 | 1.76 | 0.1876 |  |  |  |

To understand whether the occurrence of *R. saxatilis* may impact on pathogen infectivity and transmission potential, analyses were run using the *R. arcticus* and *R. saxatilis* coverages of the strain origin as explanatory variable in the model. Pathogen strain infectivity was negatively correlated with *R. saxatilis* coverage (Table 3; Fig. 4c) while in the other life-history traits, sporulation abundance, and speed to sporulation, there were no correlations (Table 3). The results of the other model variables remained qualitatively like those in the models without *R. arcticus* and *R. saxatilis* coverage (Table 3).

**Figure 4.**
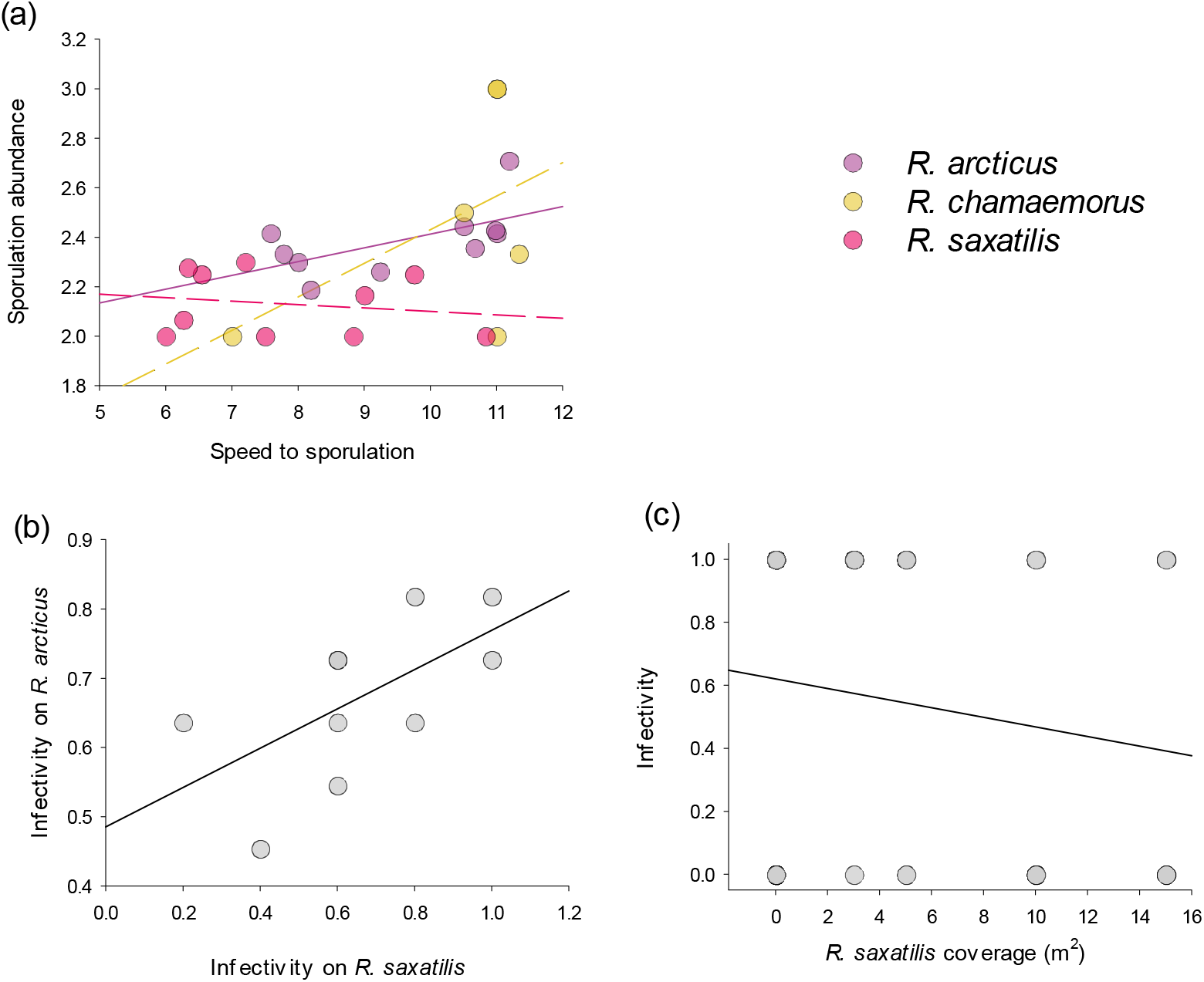
*Peronospora sparsa* life history trait correlations in three host plants in an inoculation experiment. (a) Correlation between speed to sporulation and sporulation abundance in *Rubus arcticus, R. chamaemorus*, and *R. saxatilis*. (b) Correlation between *P. sparsa* strain infectivity among the host species. (c) Correlation between *R. arcticus* population size and *P. sparsa* infection severity.

Potential life-history trade-offs limiting pathogen evolution among host species were tested by regressing the infectivity between species pairs of the three host species. There was a significant positive correlation between infectivity in *R. arcticus* and *R. saxatilis* (*P* = 0.036; *R^2^* = 0.43; Fig 4b). Correlations among *R. chamaemorus* with *R. arcticus* (*P* = 0.1268; *R^2^* = 0.27) and *R. saxatilis* (*P* = 0.3528; *R^2^* = 0.11) were non-significant. As an alternative approach to understand pathogen life history evolution among host species, I tested a correlation between speed to sporulation and sporulation abundance within each host species. I found that the sporulation abundance was explained by both host species and host genotype within species as well as speed to sporulation (Table 3; Fig. 4a). The relationship between sporulation abundance was mediated by host species (Table 3; Fig. 4a), and the positive correlation between fast and abundant sporulation observed in *R. arcticus* did not exist in *R. saxatilis* and *R. chamaemorus*.

## Discussion

Despite the commonness of generalism among pathogens and the risks of emergence of new generalist pathogens due to increased wild-life encounters, their evolution and epidemiology has gained surprisingly limited attention. Here, I investigated infectivity, transmission, and potential life-history trade-offs of *Peronospora sparsa* in its three host species *Rubus arcticus, R. chamaemorus* and *R. saxatilis* across Finland*. Rubus saxatilis* was confirmed with PCR test a novel reservoir host for this pathogen with potential to alter its epidemiology. In an epidemiological survey, I found that the disease occurrence and severity on the three host species was highest in *R. chamaemorus* and lowest in *R. saxatilis*. Disease severity varied between wild and cultivated sites. Furthermore, *P. sparsa* epidemics varied by host species composition and generally disease severity was lower in the presence of alternative host. This is interesting because the population size of each host species increased disease severity suggesting that there may be evolutionary constrains in the generalist pathogen to spread in heterogenous population (Saikkonen et al., 2016, Papaix et al., 2015). Results from a transplant experiment further showed that the host species combination in population shapes transmission dynamics of *P. sparsa*. Results from an inoculation experiment confirmed that all three plants are susceptible for the pathogen and the variation in susceptibility to *P. sparsa* is distributed among host genotypes rather than host species. Furthermore, positive life history correlations found in *R. arcticus* were altered in *R. saxatilis* suggesting a possible mechanism of evolutionary constraints (Saikkonen et al., 2016, Papaix et al., 2015).

A better understanding of life-history traits and the reservoir hosts of pathogens is needed to predict and prevent the emergence of pathogens at the interface of wild and cultivated (Morris et al., 2022). While it is known that the pathogens may freely move across wild and domesticated hosts and even across species borders (Burdon et al., 2006), the data on generalist pathogen epidemics has remained scarce. Understanding how pathogen life-history allocations are altered at the host species boundaries will help us to understand the limiting factors for the emergence of novel highly virulent strains (Papaix et al., 2015). Most of the research thus far has focused on the spill over of pathogens from cultivated plants to wild plants (Power & Mitchell, 2004). Wild plants serving as reservoir hosts for agricultural pathogens have reported amplification of epidemics and enhanced overwintering survival (Power & Mitchell, 2004). Here, I found that the symptom occurrence among wild and cultivated sites did not differ, but symptoms were more severe in the wild sites in *Rubus chamaemorus*. This is in contrast with the expectation that wild host populations are better adapted to their pathogens (Burdon & Thrall, 2009). One possible explanation for the observed pattern is that the plants in the plantations are replaced regularly with pathogen free material and thus, the disease burden will remain lighter.

Plant community composition was linked to transmission dynamics in the transplant experiment across 20 locations by comparing locations consisting of different host combinations and plant diversities. The trap plants became more often infected and had more infected leaves with increasing *R. saxatilis* coverage. However, in sites with *R. saxatilis* plant diversity decreased infection prevalence. This is in line with the expectations of dilution hypothesis and observations in other plant systems (Haas et al., 2011, Susi & Laine, 2021, Liu et al., 2020). In a study by Haas et al. (Haas et al., 2011) on an oomycete pathogen *Phytophthora ramorum* only two competent host species contributed to epidemics despite the pathogen’s wide host range. Previous studies suggest that *P. sparsa* occurs as two subspecies: one infecting hosts in genus *Rosa* and the other infecting hosts in genus *Rubus*. (Lindqvist-Kreuze et al., 2003, Koponen et al., 2000). An alternative explanation for disease variation in different plant community compositions is that in a high-quality environment the hosts may exhibit more severe symptoms (Kniskern & Rausher, 2006). The sites with higher *Rubus* plant coverage may also exhibit more favorable conditions for *P. sparsa*. However, there were no correlations between the other site explanatory factors that could explain the observed pattern. The results suggest that maintaining biodiversity is important in preventing transmission of generalist pathogens.

Warming climate may force species to shift their ranges to cooler areas (Bebber et al., 2013). Due to the global climate change, species interactions are predicted to become more intensive in high latitudes at the cool range edges (Paquette & Hargreaves, Zvereva & Kozlov, 2021). Ecological processes determining species interactions may also operate in different magnitudes along latitudinal gradients (Magnusson et al., 2020). To understand how epidemiology of the pathogen may change across latitudes, the transplant experiment and pathogen sampling for inoculation experiment were performed along a 1200 km latitudinal gradient. There was no significant linkage between latitude and symptom occurrence. However, the latitude had small but significant negative correlation with symptom prevalence in the trap plants. This is in contrast with the prediction that disease pressure would increase towards high latitudes (Bebber et al., 2013). In the inoculation experiment, the pathogens of northern origin were more infectious than southern, which is in line with the expectation of more intense interaction among species in high latitudes (Zvereva & Kozlov, 2021).

The strong effect of host genotype in the inoculation experiment as well as in the trap plant experiment was not surprising given that host related factors are expected to be in a key role in the infection outcome (Sallinen et al., 2020, Francl, 2001). The pathogen strains differed both qualitatively i.e., with different infectivity profiles, and quantitatively i.e., variation in the number of host genotypes infected. However, they did not differ in traits related to transmission potential. While there was a significant effect of latitude on the infectivity, the lack of host genotype × latitude or host genotype × latitude interactions suggests that that the resistance to *P. sparsa* may not be highly specific. Symptom occurrence and prevalence were highest in *R. chamaemorus* in field sites. However, in the inoculation experiment, two of the four genotypes *R. chamaemorus* were resistant to all *P. sparsa* strains. There are at least two possible explanations for this pattern. First, the higher disease pressure in the *R. chamaemorus* populations may lead the evolution of resistance in this species. Secondly, the pathogen strains used in the inoculation experiment originated from *R. arcticus*. It is possible that the strains naturally infecting *R. chamaemorus* may be different than those infecting *R. arcticus*.

Trade-offs are expected to limit the evolution and spread of generalist pathogens. In this study, I found that high infectivity in *R. saxatilis* and *R. arcticus* correlated positively. This suggest that a trade-off does not limit the infectivity of the pathogen between species. Instead, the strains originating from sites with high *R. saxatilis* coverage were less infectious that strains originating from sites without *R. saxatilis*. This suggests that the pathogen may still have a cost for survival in heterogeneous host populations in ability to infect a wider host range (Combes, 1997). However, there were no correlation for infectivity in *R. chamaemorus* in the infectivity in neither *R. arcticus* nor *R. saxatilis*. The three *Rubus* taxa grow commonly in shared habitats allowing the pathogens to circulate in all of them. On the other hand, the positive correlation between infectivity in *R. saxatilis* and *R. arcticus* suggests similarity of resistance in these two closely related species but difference from *R. chamaemorus*. Thus, an interesting open question for future research is to explore the genetic basis of resistance in these two host species.

Life-history trade-offs may operate in pathogen traits important for transmission to hinder the spread of the pathogen in alternative hosts (Saikkonen et al., 2016, Papaix et al., 2015). There were no significant correlations in neither speed to sporulation nor abundance among the host plants. Instead, when correlation between these two life-history traits was tested within each host species, positive correlation was found in *R. arcticus* but not in *R. saxatilis* nor *R. chamaemorus*. All the *P. sparsa* strains used in the study originated from *R. arcticus*. This result suggests that fast sporulation does not lead to high sporulation abundance in alternative hosts. This may be one mechanism of how epidemics may be hindered in populations with alternative hosts as it was seen in the survey on wild and cultivated hosts.

Pathogens may exist unnoticed in wild plants due to their mild symptoms (Prendeville et al., 2012). However, surveying the presence of the symptoms and the microbiota in wild plants is an essential approach for understanding the epidemiology and evolution of pathogen emergence. Linking a pathogen’s host range, distribution, and realized symptom severity will allow predicting the risk of disease emergence. These results shed light on the resistance variation among host species. By showing that the previously unknown reservoir hosts may have significant impact on epidemiology and evolution of a multi-host pathogen this study increases our understanding on the epidemiology of infectious diseases. When attaining novel crops into cultivation, potential risks arising from the related host species should be monitored and evaluated systematically. With the rise of novel sequencing methodologies that allow screening the microbial taxa particular attention should be paid to screening wild host microbiota for identification of the present disease risks.

## Acknowledgements

I thank Krista Raveala and Airi Lamminmäki for the maintenance of the plant and pathogen material. I am thankful for Sara Leino, Aura Palonen, and Magdalena Lukaszewicz for their help in field surveys and running the experiments. Marijke Iso-Kokkila, Eduardo Gomez, and Oskari Lehtinen are thanked for their help in the laboratory. Suvi Sallinen is thanked for helpful comments on the draft manuscript. This work was funded by the Academy of Finland (grant no 312441 to HS) and Maiju ja Yrjö Rikalan Puutarhasäätiö.

