## Supplementary Table 1 for "Alternative host shapes transmission and life-history trait correlations in a multi-host plant pathogen"

**Supplementary Material**

**Supplementary Table 1**. The sites surveyed for downy mildew *Peronospora sparsa* disease severity in *Rubus arcticus, R. chamaemorus,* and *R. saxatilis* in 2019. Mean infection prevalence and severity are shown**.**

| Site | Abbreviation | Latitude | Longitude | Species present | Infection prevalence | Mean disease severity |
|---|---|---|---|---|---|---|
| Alajärvi | ala | 62.999923 | 23.815814 | *R. arcticus* | 0.10 | 0.10 |
| Kauhajoki | kauh | 63.186563 | 23.294037 | *R. arcticus* | 0.66 | 0.06 |
| Kajaani | arm | 64.174243 | 27.715207 | *R. chamaemorus* | 0.90 | 0.33 |
| Takkarannantie, Kajaani | tak | 64.247510 | 27.839014 | *R. arcticus* | 0.50 | 0.04 |
| Takkarannantie, Kajaani | tak | 64.247510 | 27.839014 | *R. chamaemorus* | 0.80 | 0.25 |
| Honkalampi | hon | 63.025314 | 28.1729245 | *R. arcticus* | 0.50 | 0.04 |
| Honkalampi | hon | 63.025314 | 28.1729245 | *R. saxatilis* | 0.10 | 0.03 |
| Jormua | jor | 64.297225 | 27.9876965 | *R. arcticus* | 0.15 | 0.04 |
| Jormua | jor | 64.297225 | 27.9876965 | *R. saxatilis* | 0.05 | 0.02 |
| Kaurastensuo | kau | 61.031624 | 24.976035 | *R. arcticus* | 1.00 | 0.05 |
| Kaurastensuo | kau | 61.031624 | 24.976035 | *R. saxatilis* | 0.90 | 0.03 |
| Kestilä | kes | 64.183833 | 26.3703685 | *R. arcticus* | 0.90 | 0.05 |
| Kestilä | kes | 64.183833 | 26.3703685 | *R. saxatilis* | 0.90 | 0.04 |
| Koskitie, Kestilä | kos | 64.183833 | 26.370385 | *R. arcticus* | 0.30 | 0.02 |
| Koskitie, Kestilä | kos | 64.183833 | 26.370385 | *R. saxatilis* | 0.20 | 0.02 |
| Kahilantie, Kurikka | kahi | 62.567315 | 22.554436 | *R. arcticus* | 0.20 | 0.05 |
| Kahilantie, Kurikka | kahi | 62.567315 | 22.554436 | *R. saxatilis* | 0.05 | 0.02 |
| Pättikankaantie, Kurikka | pat | 62.653455 | 22.3201875 | *R. arcticus* | 0.80 | 0.08 |
| Peuraniemi | pn | 64.252572 | 27.851174 | *R. arcticus* | 0.30 | 0.11 |
| Peuraniemi | pn | 64.252572 | 27.851174 | *R. chamaemorus* | 0.60 | 0.16 |
| Peuravaara | pv | 64.637464 | 28.163065 | *R. arcticus* | 0.17 | 0.04 |
| Peuravaara | pv | 64.637464 | 28.163065 | *R. chamaemorus* | 0.68 | 0.11 |
| Ranta, Muuruvesi | rant | 63.010897 | 28.214126 | *R. arcticus* | 0.10 | 0.07 |
| Kuopio | kuo | 62.783636 | 27.502745 | *R. arcticus* | 0.95 | 0.12 |
| Kuopio | kuo | 62.783636 | 27.502745 | *R. saxatilis* | 0.30 | 0.01 |
| Kurikka | saar | 62.540290 | 22.058072 | *R. arcticus* | 0.30 | 0.04 |
| Kajaani | teb | 64.232523 | 27.7965815 | *R. arcticus* | 0.05 | 0.02 |
| Berry farm, Muuruvesi | marj | 63.012262 | 28.2140395 | *R. arcticus* | 1.00 | 0.11 |
| Varessäikkä | var | 64.884687 | 24.8196765 | *R. arcticus* | 1.00 | 0.09 |
| Varessäikkä | var | 64.884687 | 24.8196765 | *R. saxatilis* | 0.05 | 0.11 |
| Viinikka | vii | 63.162677 | 23.323661 | *R. arcticus* | 0.60 | 0.09 |
| Viinikka | vii | 63.162677 | 23.323661 | *R. chamaemorus* | 0.30 | 0.19 |

**Supplementary Table 2.** The sites used in the *Rubus arcticus* trap plant experiment in 2021.

| Site | Abbreviation | Latitude | Longitude | Infection prevalence | Mean disease severity |
|---|---|---|---|---|---|
| Honkalampi | HON | 63.025314 | 28.1729245 | 0.58 | 0.17 |
| Hongikonluoma | HONG | 63.177292 | 23.32972652 | 1.00 | 1.00 |
| Kaihlakyrö | KAIH | 62.508367 | 22.5961235 | 0.93 | 0.37 |
| Kestilä | KEST | 64.183833 | 26.3703685 | 0.77 | 0.19 |
| Kevo | KEV | 69.723507 | 27.0408455 | 0.83 | 0.55 |
| Station, Kevo | KEVS | 69.757128 | 27.0107795 | 0.77 | 0.34 |
| Korretoja | KOR | 69.623164 | 27.128794 | 1.00 | 0.52 |
| Kortteisen tekojärvi | KORJ | 64.174479 | 26.1048475 | 0.90 | 0.22 |
| Muuruvesi | MARJ | 63.012262 | 28.2140395 | 0.92 | 0.67 |
| Ojaniementie | OJA | 62.47891 | 22.3608345 | 0.91 | 0.81 |
| Pahkalampi | PAH | 63.1246021 | 28.17965175 | 0.55 | 0.20 |
| Pättikankaantie, Kurikka | PÄT | 62.653455 | 22.3201875 | 0.79 | 0.35 |
| Revonkanta | REV | 64.419568 | 28.4196135 | 0.50 | 0.12 |
| Ristimaantie | RIS | 63.15507 | 23.09796815 | 0.67 | 0.63 |
| Säärenperä | SÄÄ | 64.90069 | 25.01446965 | 1.00 | 0.96 |
| Takalahdenkuja | TAKA | 63.22758 | 23.26324448 | 0.84 | 0.29 |
| Tavastperä | TAV | 64.169853 | 26.4197305 | 0.96 | 0.20 |
| Kajaani | TEB | 64.232523 | 27.7965815 | 0.71 | 0.57 |
| Tikkaperä | TIK | 64.76379 | 25.2793305 | 0.91 | 0.40 |
| Varessäikkä | VAR | 64.884687 | 24.8196765 | 0.69 | 0.15 |

**Supplementary Table 3.** The material used in the *Peronospora sparsa* inoculation experiment on *Rubus arcticus, R. chamaemorus* and *R. saxatilis.*

| Material | Site of origin | latitude | longitude |
|---|---|---|---|
| *Peronospora sparsa* strains | | | |
| HONG | Hongikonluoma | 63.17729 | 23.32973 |
| KAIH | Kaihlakyrö | 62.50837 | 22.59612 |
| KEV | Kevo | 69.72351 | 27.04085 |
| MARJ | Berry farm, Muuruvesi | 63.01226 | 28.21404 |
| OJA | Ojaniementie | 62.47891 | 22.36083 |
| PAH | Pahkalampi | 63.1246 | 28.17965 |
| SÄÄ | Säärenperä | 64.90069 | 25.01447 |
| TAV | Tavastsäikkä | 64.16985 | 26.41973 |
| TEB | Kajaani | 64.23252 | 27.79658 |
| TIK | Tikkaperä | 64.76379 | 25.27933 |
| *Rubus arcticus* genotypes | | | |
| Pima | Cultivar | | |
| kos | Koskitie, Kestilä | 64.183833 | 26.370385 |
| P | Pättikankaantie, Kurikka | 62.65346 | 22.32019 |
| Teb | Kajaani | 64.23252 | 27.79658 |
| kau | Kaurastensuo | 61.031624 | 24.976035 |
| KEST | Kestilä | 64.18383 | 26.37037 |
| m22 | Honkalampi | 63.02531 | 28.17292 |
| JOM | Jormua | 64.29723 | 27.98769 |
| HONG | Hogikonluoma | 63.17729 | 23.32973 |
| VAR | Varessäikkä | 64.88469 | 24.81968 |
| VII | Viinikka | 63.162677 | 23.323661 |
| *Rubus chamaemorus* genotypes | | | |
| Haaga | Haaga | 60.22158 | 24.89354 |
| LAM | Lamminjärvi | 61.07532 | 25.0505 |
| VIIN | Viinikka | 63.162677 | 23.323661 |
| ARM | Kajaani | 64.174243 | 27.715207 |
| *Rubus saxatilis* genotypes | | | |
| VA | Varessäikkä | 64.88469 | 24.81968 |
| ÅI | Åland Islands | 60.16973 | 19.97117 |
| HON | Honkalapi | 63.02531 | 28.17292 |
| kes | Kestiläntie | 64.18383 | 26.37037 |
| kevo | Kevo | 69.72351 | 27.04085 |

**Supplementary Table 4.** The results from regression analyses among the site variables in the *Rubus arcticus* trap plant experiment.

| Latitude | $R^2$ | *P* |
|---|---|---|
| *R. chamaemorus* coverage | 0.0188 | 0.5582 |
| *R. saxatilis* coverage | 0.0858 | 0.1976 |
| *R. arcticus* coverage | 0.1491 | 0.0921 |
| *Shannon's* diversity | 0.0321 | 0.4495 |

| *Shannon's* diversity | $R^2$ | *P* |
|---|---|---|
| *R. chamaemorus* coverage | 0.142 | 0.1015 |
| *R. saxatilis* coverage | 0.061 | 0.2938 |
| *R. arcticus* coverage | 0.002 | 0.9564 |

| *R. arcticus* coverage | $R^2$ | *P* |
|---|---|---|
| *R. chamaemorus* coverage | 0.0816 | 0.2094 |
| *R. saxatilis* coverage | 0.0039 | 0.7868 |

**Supplementary Figure 1.** (a) The wild and cultivated sampling and survey locations of *Peronospora sparsa* on *Rubus arcticus, R. chamaemorus,* and *R. saxatilis* in 2019. Symptoms of *P. sparsa* on (b) *R. arcticus*, (c) *R. chamaemorus*, (d) *R. saxatilis*.

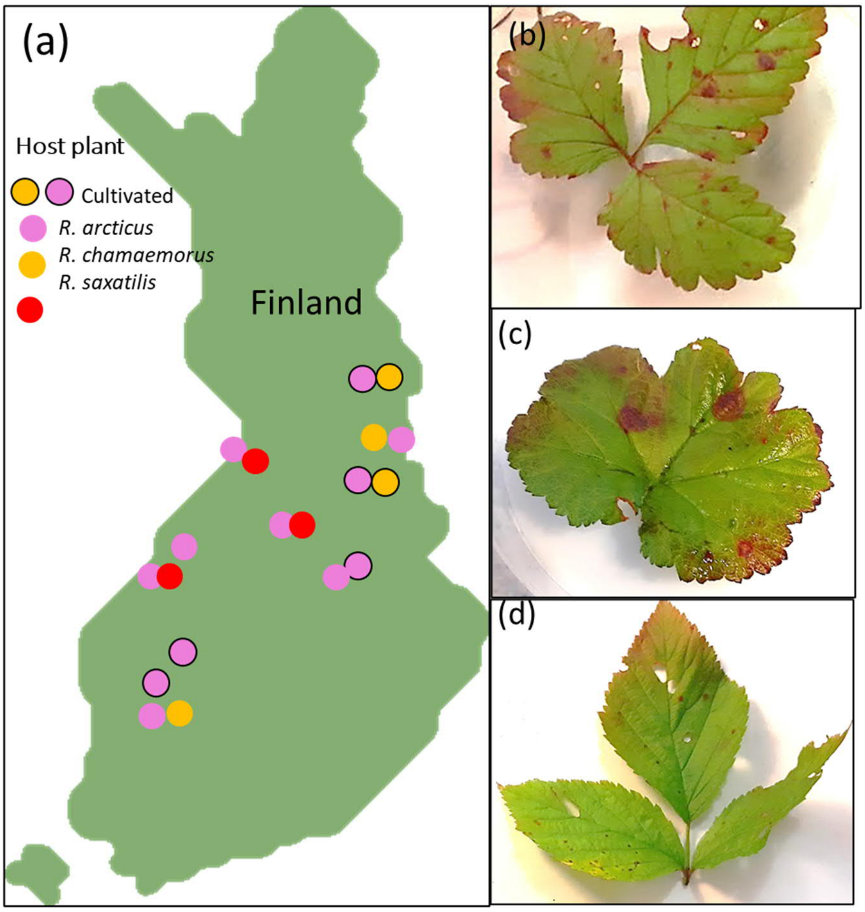
